# Origin of perseveration in the trade-off between reward and complexity

**DOI:** 10.1101/2020.01.16.903476

**Authors:** Samuel J. Gershman

## Abstract

When humans and other animals make repeated choices, they tend to repeat previously chosen actions independently of their reward history. This paper locates the origin of perseveration in a trade-off between two computational goals: maximizing rewards and minimizing the complexity of the action policy. We develop an information-theoretic formalization of policy complexity and show how optimizing the trade-off leads to perseveration. Analysis of two data sets reveals that people attain close to optimal trade-offs. Parameter estimation and model comparison supports the claim that perseveration quantitatively agrees with the theoretically predicted functional form.

## Introduction

In his pioneering work on animal intelligence, Thorndike (1911) formulated several “laws” of learning. The most famous of these, the *law of effect*, stated that an action yielding a reward will become more likely to be repeated in the future. The lesser-known *law of exercise* stated that simply producing an action will make it more likely to be repeated in the future (and concomitantly, an action will become less likely to be repeated if it’s not produced). The law of exercise implies a form of perseveration: with sufficient frequency of repetition, an action will be selected even if it no longer yields the highest reward among the possible choices. The law of exercise thus captures a key signature of habit, whereby repetition causes behavior to become “autonomous” from the agent’s goals (Dickinson, 1985; Miller et al., 2019; Wood and Rünger, 2016).

Although these laws do not exhaustively determine action selection, they are supported by many studies of humans and other animals. For example, Lau and Glimcher (2005) found that monkeys performing a two-alternative forced choice task were influenced by the recent history of both rewards and choices, a finding that also extends to human subjects (e.g., Seymour et al., 2012). Under time pressure, people will frequently repeat previous actions despite intending to choose an alternative action (Betsch et al., 2004). Reaction times are also facilitated for response repetitions in serial choice reaction tasks (e.g., Bertelson, 1965; Rabbitt and Vyas, 1974). In everyday life, past actions predicts future actions (e.g., product choices in the supermarket; Riefer et al., 2017), even after controlling for other predictors such as conscious intentions and beliefs about social norms (Ouellette and Wood, 1998).

While perseveration has been ubiquitously documented, a basic puzzle is *why* it occurs at all. If the goal is to maximize reward, an agent’s actions should be entirely predictable from its reward history; in other words, the law of exercise should be completely dominated by the law of effect. If anything, the need to explore actions in order to gain information about their consequences should induce a tendency *against* repeating past actions (Riefer et al., 2017; Schulz and Gershman, 2019).

A common theme in the psychology of habit is the idea that perseveration is somehow less effortful (Wood and Rünger, 2016). Some reinforcement learning models have conceptualized effort in terms of computational complexity; habits arise from action selection based on a look-up table of cached reward expectations, which demands less effort compared to goal-directed action selection based on planning with an internal model of the task (Daw, 2018). Consistent with this conceptualization, taxing cognitive resources (for example, by having subjects perform a secondary task or increasing the difficulty of planning) results in greater reliance on habit (Gershman et al., 2014; Kool et al., 2018b; Otto et al., 2013a). The problem with this view of habit, as pointed out by Miller et al. (2019), is that it does not exactly correspond to Thorndike’s law of exercise: caching reward expectations in a look-up table does not by itself produce a bias to repeat actions. Rather, this form of caching can be viewed as implementing Thorndike’s law of effect.

Miller et al. (2019) propose an alternative model that explicitly formalizes the law of exercise, whereby taking an action increases its habit strength independently from reward. While this model succeeds as a descriptive account of habitual action selection, it does not provide a computational rationale for perseveration. Thinking about this rationale in terms of computational complexity seems unpromising, since it’s not obvious why looking up a cached habit value would be less cognitively expensive than looking up a cached reward expectation, and the latter is obviously more useful from the perspective of reward maximization.

A different approach to this question rests upon a distinction between *computational* and *statistical* complexity. Whereas computational (or time) complexity measures how much thinking is required to perform a task, statistical (or sample) complexity measures how much learning is required. Effort, in this case, corresponds to the difficulty of learning. Is it possible that habits are less statistically complex? In fact, Filipowicz et al. (2020) have shown that learning cached reward expectations is not necessarily more statistically complex than learning an internal model for planning (they did not directly address the perseverative notion of habit).

In this paper, we explore a different computational rationale for perseveration, based on the notion of *policy* complexity (Lerch and Sims, 2018; McNamee et al., 2016; Parush et al., 2011; Still and Precup, 2012; Tishby and Polani, 2011). In the language of reinforcement learning theory, a policy *π*(*α|s*) is a probabilistic mapping from states to actions (Sutton and Barto, 2018), where a state corresponds to the information about the environment that is needed for reward prediction. To implement a policy computationally, we would need to describe it in some programming language, and the description length of that program (e.g., in bits or nats) imposes a demand on memory resources. Intuitively, if a policy can be “compressed” to a short description length, it will be easier to remember, much in the same way that the benefits of compression have been studied in memory for symbolic and visual stimuli (Brady et al., 2009; Mathy and Feldman, 2012; Nassar et al., 2018). As we will formalize later, it turns out that perseveration arises naturally from the imperative to reduce policy complexity. Perseveration is, in essence, a form of policy compression.

Policy complexity is conceptually different from computational complexity; one could have a policy with low policy complexity and high computational complexity, or vice versa. For example, finding the shortest path between two distant cities might require an expensive optimization (high computational complexity), but the optimal path itself might be very simple, like staying on one highway for most of the trip (low policy complexity). In contrast, finding the shortest path between two locations in the same city might be cheap (low computational complexity), but the optimal path might be tortuous (high policy complexity), as anyone who has tried to get around Boston by car knows well.

The question addressed here is how people negotiate the trade-off between reward and policy complexity. The mathematical toolbox for answering this question comes from the branch of information theory known as *rate distortion theory* (Berger, 1971). The next section reviews the elementary concepts as they apply to policy optimization. Rate distortion theory allows us to derive the optimal trade-off function, which reveals that perseveration will occur for any resource-bounded agent. Since both reward and policy complexity are experimentally measurable, we can evaluate the degree to which choice data conform to the optimal trade-off function. Furthermore, by fitting parametrized policies to the choice data, we can evaluate how well the data match the theoretically predicted form of perseveration.

## Methods

All code and data for reproducing the analyses described below is available at https://github.com/sjgershm/reward-complexity.

### Theoretical framework

Rate distortion theory addresses the interface between information theory and statistical decision theory. Here we will adopt somewhat non-standard terminology, following Parush et al. (2011), in order to draw a clearer connection with the issues raised in the Introduction. We will assume that an agent either learns or has direct access to a *value function Q*(*s, a*) that defines the expected reward in state *s* after taking action *a*. Each state is visited with probability *P*(*s*), and an action is chosen according to a policy *π*(*α|s*). In the language of information theory, we can think of the state distribution as a *source* and the policy as a *noisy channel*, mapping messages (states) to codewords (the internal representation), which are in turn mapped to output signals (actions). The average codeword length (or *rate*) necessary to encode a policy with arbitrarily small error is equal to the mutual information between states and actions:

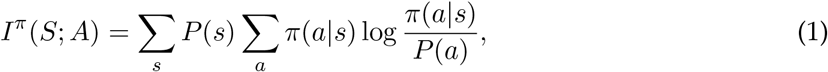

where *P*(*a*) Σ_*s*_ *P*(*s*)*π*(*α|s*) is the marginal probability of choosing action *a* (i.e., the policy averaged across states). Because the mutual information quantifies the degree of probabilistic dependency between states and actions, we will refer to it as the *policy complexity*. Intuitively, policies are more complex to the extent that the policy is state-dependent. If the policy is the same in every state, then the policy complexity is minimized (mutual information is equal to 0).

The communication channel formulation is useful because it lets us see why compression makes sense. Real-world environments involve an astronomical number of states and actions, so a resource-limited system can’t afford to represent all of them in a giant look-up table. This goes against the conventional wisdom that look-up tables are computationally cheap (e.g., Kool et al., 2018a); although they require little thinking (low computational complexity), they require a large number of bits (high policy complexity). Previous applications of rate distortion theory to psychology have used this insight to explain the factors influencing confusability in memory, on the assumption that items cannot be stored in a look-up table due to resource constraints (Sims et al., 2012; Sims, 2016).

Exactly how much to compress depends on the amount of reward that can be achieved for a given policy complexity. Let us denote the average reward by:

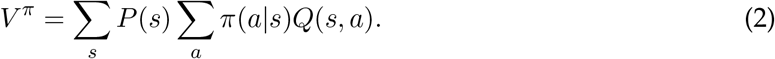

We can now formulate the optimization problem:

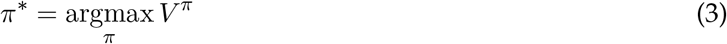

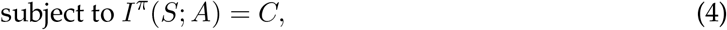

where *C* denotes the channel capacity—the maximum achievable policy complexity. Intuitively, the goal is to earn as much reward as possible, subject to the constraint that the policy complexity cannot exceed the capacity limit. We have left implicit two other necessary constraints (action probabilities must be non-negative and sum to 1). This constrained optimization problem can be rewritten in a Lagrangian form:

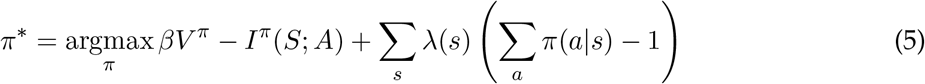

with Lagrange multipliers *β* and λ(*s*). The optimal policy *π*\* has the following form (Parush et al., 2011; Still and Precup, 2012; Tishby and Polani, 2011):

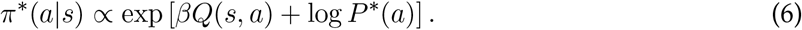

A number of recognizable components now come into view. The optimal policy is a softmax function, used ubiquitously in the reinforcement learning literature for both simulating artificial agents and modeling biological agents. The Lagrange multiplier *β* plays the role of the “inverse temperature” parameter, which regulates the exploration-exploitation trade-off via the amount of stochasticity in the policy (Sutton and Barto, 2018). When *β* is close to 0, the policy will be near-uniform, and as *β* increases, the policy will become increasingly concentrated on the action with maximum value. However, note that the derivation of the optimal policy makes no reference to exploration (see Still and Precup, 2012). Rather, *β* reflects the resource constraint—more precisely, its inverse is the partial derivative of the value with respect to the policy complexity:

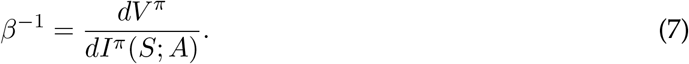

This means that the inverse temperature for the optimal policy will be lower (choice stochasticity higher) when varying the policy complexity has a greater effect on reward (see also Ortega and Braun, 2013; Zénon et al., 2019).

Another important property of Eq. 6 is the log *P*\* (*a*) term, which arises from the need for policy compression due to the capacity constraint. This implies that frequently chosen actions should bias the policy (i.e., produce perseveration), in accordance with Thorndike’s law of exercise. We will empirically evaluate the specific functional form of perseveration implemented by Eq. 6, as described below.

The perseveration term implicitly depends on the optimal policy, since

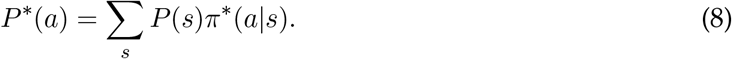

Thus, to find the optimal policy, we can use a variation of the classic *Blahut-Arimoto algorithm* (Arimoto, 1972; Blahut, 1972), alternating between updating the policy according to Eq. 6 and updating the marginal action distribution according to Eq. 8. By performing this optimization for a range of *β* values, we can construct a *reward-complexity curve* that characterizes the optimal policy for a given resource constraint. That is, for a given resource constraint, the point on the reward-complexity curve yields the highest reward with the least amount of perseveration. The slope of each point on the reward-complexity curve is given by *β*^-1^. The reward-complexity curve is always concave, which means that *β* monotonically increases with policy complexity.

### Why mutual information measures perseveration

Although not immediately obvious, the mutual information between states and actions provides an intuitive measure of perseveration. Consider an agent that adopts the same action policy regardless of what state it’s in. Mathematically, this implies that 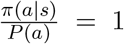, or equivalently as 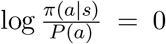. This policy produces a high-level of perseveration, because an agent will tend to continue taking the same actions even after the state has changed. If the agent adopts a statedependent policy, and hence perseverates less, then the log probability ratio will on average be greater than 0. Thus, the average log probability ratio is monotonically related to the degree of perseveration. This quantity is in fact just the mutual information. It is lower-bounded by 0 (maximum perseveration) and upper-bounded by the entropy of the marginal action distribution 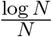, where *N* is the number of actions. This upper bound is achieved by an agent that deterministically selects actions in each state such that the marginal distribution over actions is uniform.

It is important to note that mutual information is a theory-agnostic measure of perseveration in the sense that it makes no assumption about how agents negotiate the reward-complexity trade-off, or indeed about how they make decisions at all. Thus, although it is identical to policy complexity (a theory-based concept), we can always interpret the complexity axis of reward-complexity plots as a measure of perseveration.

### Data sets

We evaluated the predictions of the theory developed in the previous section using two data sets. The first data set, reported in Collins (2018), consists of 91 subjects performing a reinforcement learning task in which the set size (the number of distinct stimuli, corresponding to states) varied across blocks (Figure 1A). On each trial, subjects saw a single stimulus, chose an action and received reward feedback. Each stimulus was associated with a single rewarded action. The experiment consisted of a learning and test phase (with no reward feedback), but here we only analyze the learning phase data. Each subject completed 14 blocks, half with set size 3 and half with set size 6. Each stimulus appeared 12-14 times in a block. No stimulus was repeated across blocks.

**Figure 1:**
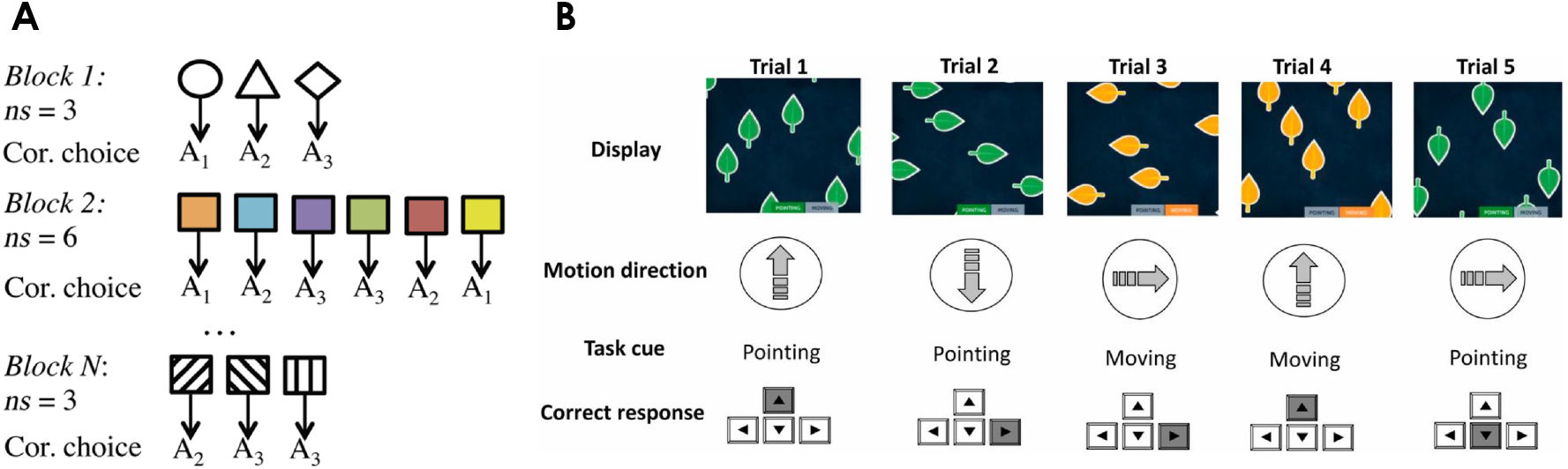
Schematic of experimental tasks. (A) In Collins (2018), subjects saw a single stimulus on each trial and then chose between 3 actions. Each stimulus corresponded to a state with a single rewarded action. The number of stimuli was varied across blocks. (B) In Steyvers et al. (2019), subjects viewed a display containing leaves moving coherently in one of 4 cardinal directions. The leaves also pointed in one of four cardinal directions. On some trials (indicated by orange leaf color) subjects made a motion direction judgment, and on other trials (indicated by green leaf color) subjects made a pointing direction judgment. Feedback was provided after each judgment.

The second data set, reported in Steyvers et al. (2019), consists of 1000 subjects playing the task-switching game “Ebb and flow” on the Lumosity platform (Figure 1B). On each trial, subjects viewed moving leaves on a display and reported either the motion or pointing direction of the leaves. In this case, the state corresponds to a tuple (task, motion direction, pointing direction), defining 32 distinct states. Subjects played between 371 and 5227 trials, with a median of 2735 trials (99% of subjects played over 1000 trials, so the task can be considered well-practiced for most subjects).

### Mutual information estimation

To construct the empirical reward-complexity curve, one needs to estimate two quantities: average reward and the mutual information between states and actions. Estimation of average reward is straightforward, but estimation of mutual information is notoriously tricky (see Paninski, 2003). We used the Hutter estimator, which computes the posterior expected value of the mutual information under a Dirichlet prior (Hutter, 2002). We chose a symmetric Dirichlet prior with a concentration parameter *α* = 0.1, which exhibits reasonably good performance when the joint distribution is sparse (Archer et al., 2013).^1^ The sparsity assumption is likely to hold true in the data sets analyzed here because there is a single rewarded action in each state. As shown in the Results, this produced empirical reward-complexity curves that mostly satisfied the theoretical bound.

### Parameter estimation and model comparison

To quantitatively evaluate the theory, we fit models of the following form:

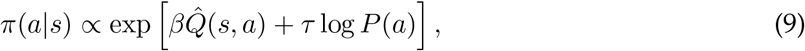

where 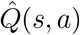 is an estimate of the expected reward and *τ* is a choice perseveration parameter. In model M1, we fit both *β* and *τ* as free parameters. Previous models incorporating a perseveration parameter typically treated it as a purely descriptive device, soaking up a large source of variance (e.g., Gershman, 2016; Lau and Glimcher, 2005; Seymour et al., 2012). These earlier models did not typically place constraints on the parameter value, and nor have we in this paper. Critically, the rate distortion framework *does* make predictions about the parameter value, namely that it should equal 1 when *β* is allowed to vary. Accordingly, in model M2 we fit only *β*, and forced *τ* to equal 1 (corresponding to the optimal policy in Eq. 6). Maximum likelihood parameter estimates were obtained using unconstrained optimization with 5 random initializations to avoid local maxima.

Model comparison was performed using a Bayesian random effects procedure (Rigoux et al., 2014). In brief, this procedure estimates the population-level frequency of each model, along with the probability that an individual subject’s data were generated by each model. We report the log model evidence favoring M2 over M1 for each subject, as well as the protected exceedance probability, which measures the probability that M2 is more likely in the population than M1, taking into account the probability of spurious differences due to randomness.

The procedure to obtain 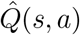 and *P*(*a*) for each trial was slightly different for the two data sets. For the Collins (2018) data set, the Q-values were initialized to 0, learning was modeled using a standard delta rule:

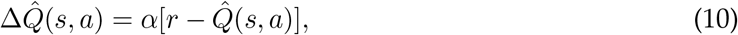

where α is a learning rate parameter (which was fitted to the data) and *r* is the received reward. The marginal action probability *P*(*a*) was estimated using a 5-trial moving average (truncated at boundaries between blocks).

For the Steyvers et al. (2019) data set, we assumed that subjects had full knowledge of *Q*(*s, a*) and simply hard-coded it into the policy. As with the Collins data set, the marginal action probability was estimated using a 5-trial moving average. Since there was no discrete block structure, no truncation of the moving average was applied.

## Results

To briefly recapitulate the key points from the theoretical framework: if there are a limited number of bits available to encode a policy (the capacity constraint), then the reward-maximizing policy subject to this constraint will be *compressed*, ignoring some state information.^2^ Compression implies perseveration, in the sense that actions will be selected in proportion to their frequency of past selection (a form of Thorndike’s Law of Exercise). If the perseveration lies on the reward-complexity curve, we can describe it as achieving an optimal trade-off between reward and policy complexity under a particular capacity constraint, which may vary across individuals. The two main goals of this section are (1) to evaluate whether individuals do in fact lie near the reward-complexity curve, and (2) to evaluate whether action selection follows the specific parametric model dictated by the optimal resource-constrained policy.

Figures 2 and 3 show the reward-complexity curves for the two data sets, with the empirical data superimposed. As predicted by the theory, reward generally increases monotonically with policy complexity, with values close to the optimal trade-off curve. Figure 2 also shows that policy complexity is higher for larger set sizes, resulting in lower average reward.

**Figure 2:**
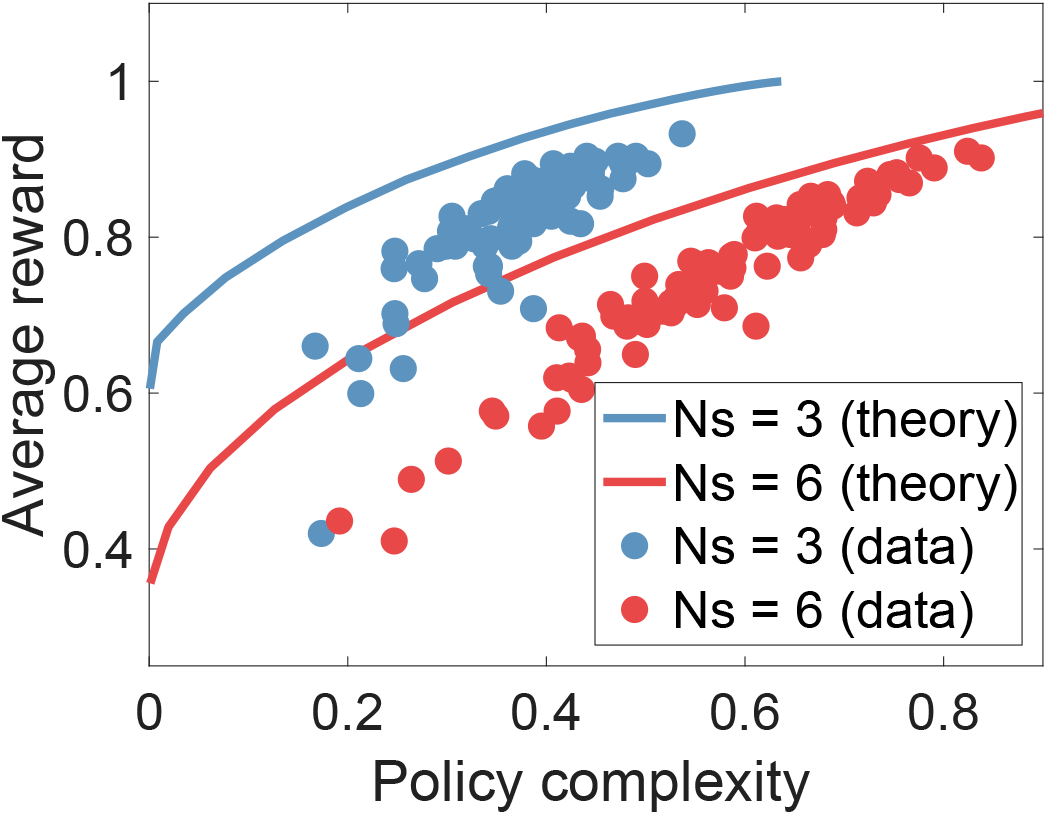
The reward-complexity trade-off, applied to data from Collins (2018). Each solid line shows the optimal trade-off function for a particular set size (Ns = 3 or 6). The circles show data from different blocks of trials, aggregated across subjects. Complexity is measured in nats.

**Figure 3:**
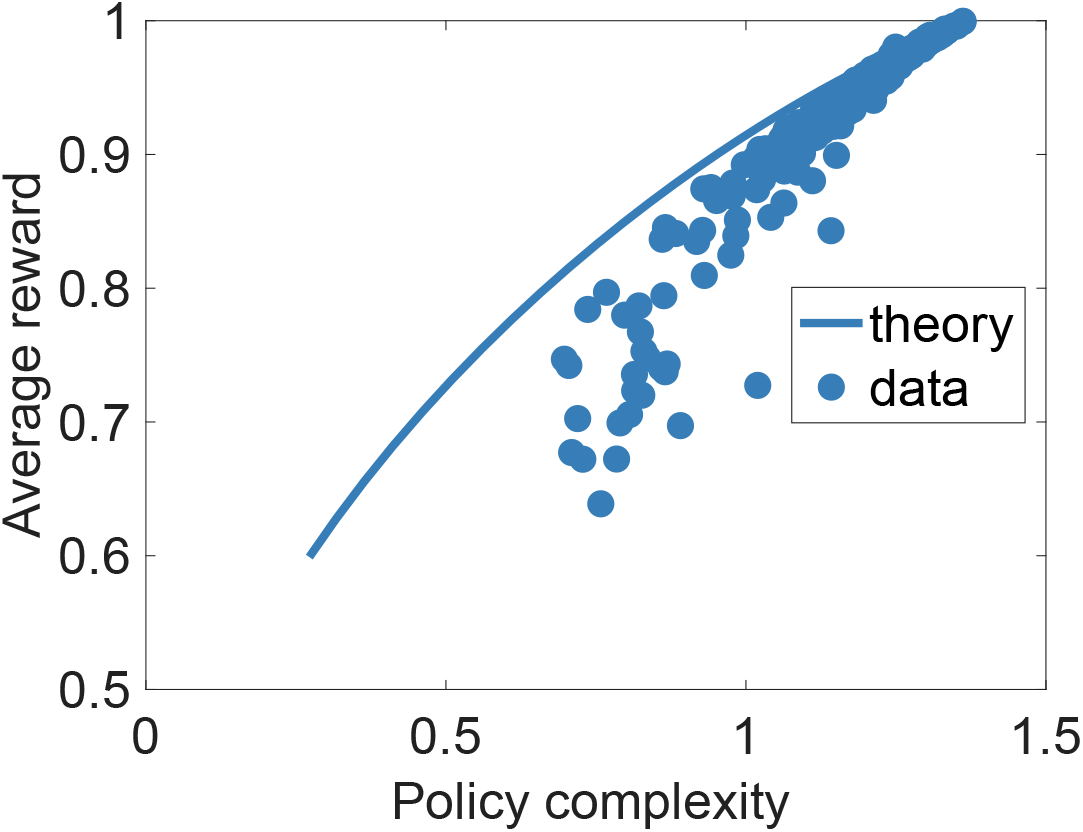
Reward-complexity trade-off, applied to data from Steyvers et al. (2019). The solid line shows the optimal trade-off function, and the circles correspond to individual subjects. Complexity is measured in nats.

To quantify the agreement between theory and data, we used interpolation to identify the predicted average reward for each measured policy complexity value. These predictions were significantly correlated with the empirical average reward (*r* = 0.91 for the Collins data set, *r* = 0.96 for the Steyvers data set, both *p* < 0.00001). Despite this quantitative agreement, the data also indicate a salient deviation from the optimal reward-complexity curve: subjects with low policy complexity achieve lower average reward than would be predicted by the optimal policy. We quantified this by computing the correlation between the bias (how far a subject is from the theoretical curve) and policy complexity, finding a significant negative correlation for both data sets each measured policy complexity value (*r* = −0.55 for the Collins data set, *r* = −0.81 for the Steyvers data set, both *p* < 0.00001).

We next sought to evaluate the functional form of the policy described by Eq. 6. As described in the Methods, we fit two models to the choice data: M1, which fits the degree of choice perseveration as a free parameter to each subject separately, and M2, which forces the parameter to equal 1 (in accordance with the theory). The distributions of estimates for this parameter are shown in the left panels of Figure 4, revealing that they are concentrated around 1 (89% of the parameter estimates were between 0.5 and 1.5 for the Collins data set, and 96% in the Steyvers data set). This conclusion was further validated by random-effects Bayesian model selection, which strongly favored model M2 over M1 (protected exceedance probability greater than 0.99 for both data sets). M2 was favored for almost all subjects, as shown in the right panels of Figure 4. Taken together, these results show that the resource-constrained optimal policy provides a good quantitative model of perseveration in these data sets.

**Figure 4:**
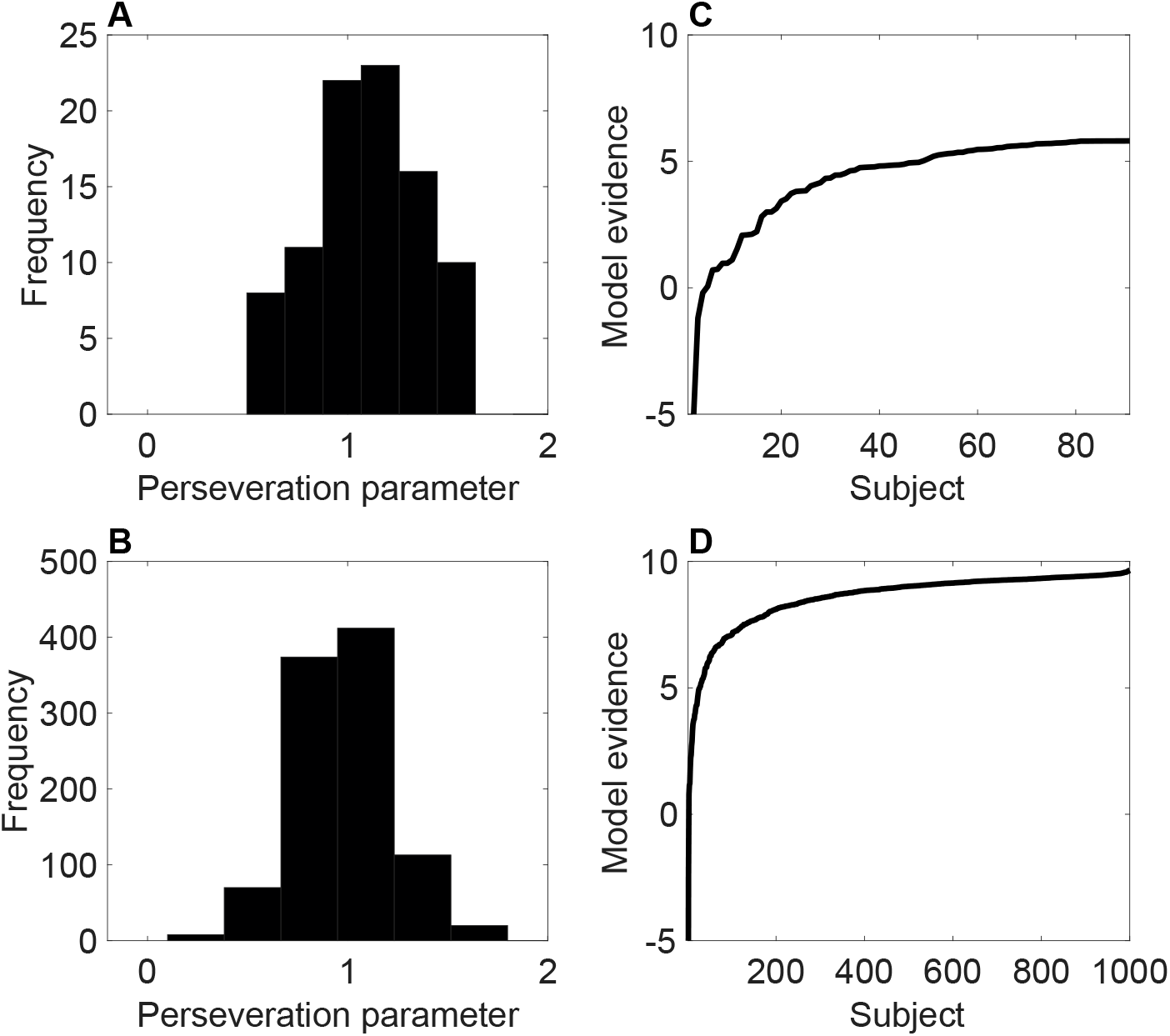
(A,B) Histogram of maximum likelihood estimates for the choice perseveration parameter. (C,D) Log posterior odds in favor of M2 (the optimal trade-off model) for each subject, sorted by increasing evidence. An evidence of 0 indicates equal support for M1 and M2. The top panels show the results for the data from Collins (2018); the bottom panels show the results for the data from Steyvers et al. (2019).

Although M2 was favored on aggregate, the relative evidence for M2 over M1 for individual subjects was negatively correlated with each subject’s bias (*r* = −0.30 for the Collins data set, *r* = −0.25 for the Steyvers data set, both *p* < 0.01). This indicates that deviations from optimality might be partly explicable in terms of Eq. 9. Indeed, the estimated *τ* parameter was negatively correlated with bias (*r* = −0.55 for the Collins data set, *r* = −0.09 for the Steyvers data set, both *p* < 0.01). Thus, subjects with higher bias tended to have higher levels of perseveration, consistent with the empirical reward-complexity curves shown in Figure 2 and 3.

To ensure that our parameter estimation results are not a spurious consequence of the model structure (e.g., due to identifiability issues), we simulated data from Eq. 9 applied to the experimental design from the Steyvers study. The inverse temperature and choice perseveration parameters were sampled uniformly from the range [0, 5]. We then fit the model to these simulated data using the same procedure that we applied to the experimental data. Figure 5 shows a tight correlation between the true and recovered choice perseveration parameter estimates (*r* = 0.94), indicating that this parameter is indeed recoverable, bolstering our confidence in the analyses of parameter estimates for the experimental data.

**Figure 5:**
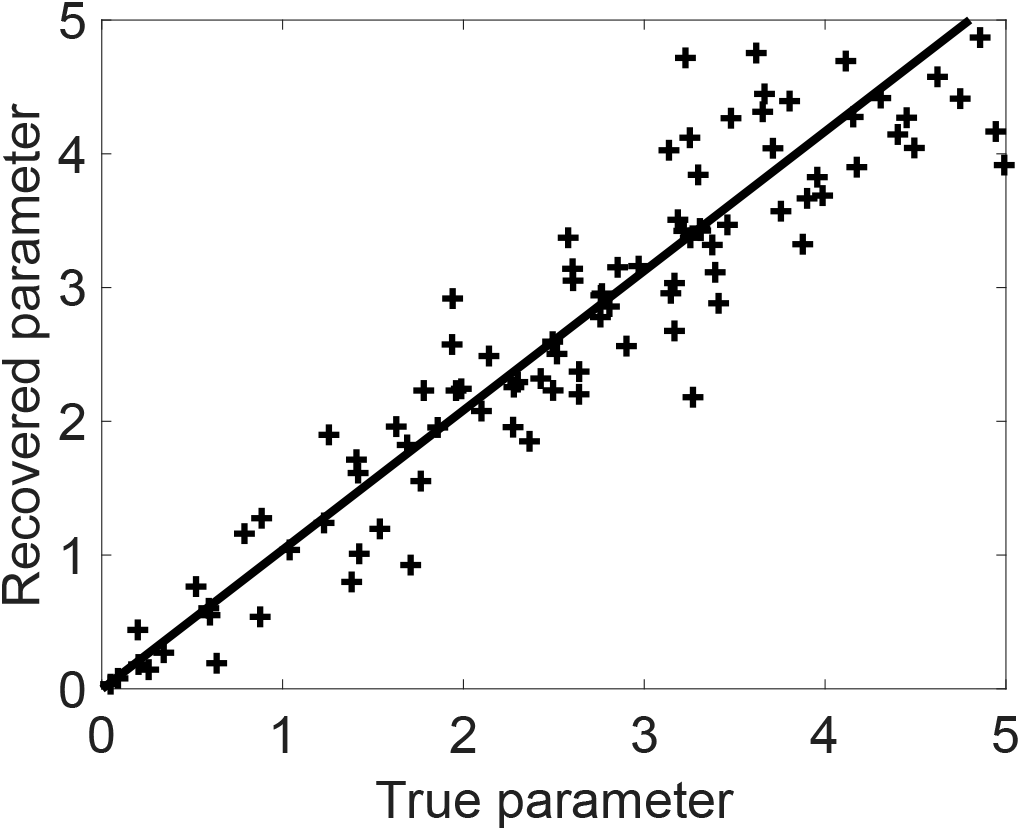
Parameter recovery from simulated data. Solid line shows the least-squares fit.

## Discussion

The idea that many aspects of cognition can be explained in terms of resource-bounded rationality has gained wide currency (Gershman et al., 2015; Lieder and Griffiths, 2019). The precise nature of the resource constraints and their implications is a matter of ongoing research. We contribute to this line of thought by applying rate distortion theory to a fundamental puzzle in psychology: why do humans and other animals perseverate? The answer provided here is that perseveration is a natural consequence of limitations on policy complexity. If the number of bits available to encode a policy is finite, then a resource-rational agent will exhibit perseveration. We showed empirically, using two data sets, that subjects or task conditions with different degrees of policy complexity yield predictable levels of reward attainment in accordance with the optimal reward-complexity trade-off. Our analyses also showed that the functional form of perseveration was quantitatively consistent with rate distortion theory. Nonetheless, there was a systematic deviation from the optimal trade-off function for subjects with low policy complexity.

Why was the deviation from optimality higher for low complexity subjects? Our data do not provide a definitive answer. One possibility is that optimization of the resource-constrained objective function is itself resource-constrained, such that people who can devote fewer bits to encoding their policy also have fewer computational resources to find the optimal solution. This would be consistent with evidence that working memory capacity predicts the deployment of computationally expensive planning algorithms (Gershman et al., 2014; Otto et al., 2013b; Schad et al., 2014). Another possibility is that low complexity subjects are not optimizing a resource-constrained objective function at all, instead relying on heuristics that are computationally cheap but sub-optimal (Gigerenzer and Gaissmaier, 2011). Teasing apart these hypotheses will require new experiments to measure individual differences in various cognitive capacities, as well as more explicit hypotheses about heuristics that quantitatively predict the deviation from optimality.

In the Introduction, we highlighted a distinction between statistical complexity (the amount of data needed to learn a policy) and policy complexity (the number of bits needed to encode a policy). However, these concepts are connected, because simpler policies are more easily learned. This follows from the general principle that *compression implies learning* (Blum and Langford, 2003), which can be formalized in a number of ways. For example, in the setting where *Q*(*s, a*) = 1 if a ∈ {0,1} is correct and 0 otherwise, the policy can be viewed as a binary classifier and the rewards can be viewed as labels (the standard supervised learning problem). Roughly speaking, if the number of bits required to describe the policy is much less than the number of samples, then we can guarantee accurate generalization to new samples (Blumer et al., 1987). The connection between compression and learning explains why the mutual information between states and actions can be used to measure both statistical complexity (see Filipowicz et al., 2020) and policy complexity.

The theoretical framework of rate distortion theory is highly abstract. We have made very few assumptions about the underlying cognitive mechanisms that produce a particular point on the reward-complexity curve. This contrasts with the modeling that was previously applied to the same data sets (Collins, 2018; Steyvers et al., 2019), which explored detailed mechanistic hypotheses. These different approaches have different advantages and disadvantages. Ultimately, we would like detailed mechanistic theories of cognition of the sort developed by Collins, Steyvers, and their colleagues. At the same time, the search for general principles can be usefully pursued at a more abstract level of the sort developed here. This has the advantage of allowing us to make general claims about the nature of cognition that transcend particular mechanistic implementations.

One important source of data for mechanistic implementations of decision making is response time. Of particular relevance is recent work by Urai et al. (2019), who studied the relationship between response time and *choice-history bias* in perceptual decision making—the robust finding that decisions are biased towards repetition across trials, even when the perceptual evidence is uncorrelated (Braun et al., 2018; Fründ et al., 2014; Howarth and Bulmer, 1956; Verplanck et al., 1952). Using a sequential sampling model, Urai and colleagues argued that choice history alters the rate of evidence accumulation, such that evidence in favor of previous choices is weighted more strongly (a form of confirmation bias; see also Abrahamyan et al., 2016; Talluri et al., 2018). One interpretation of this finding is that the locus of policy compression in perceptual decision tasks originates at the level of attention to stimulus information rather than at the level of the policy. More generally, policy compression could arise from any process along the sensory-to-motor mapping that reduces mutual information. It is a task for future work to catalogue and disentangle the effects of these processes.

Rate distortion theory holds promise as a vehicle for general principles because it unifies two frameworks (information theory and statistical decision theory) that already by themselves have broad explanatory reach. Rate distortion theory has been successfully applied to many different cognitive phenomena, ranging from working memory (Sims et al., 2012; Sims, 2016) and absolute identification (Sims, 2018) to language (Zaslavsky et al., 2018) and motor control (Schach et al., 2018). A complete theory in these domains will eventually use mechanistic models to constrain the rate distortion analysis.

## Acknowledgments

I am grateful to Anne Collins and Mark Steyvers for making their data available, and to Rahul Bhui, Rani Moran, Lucy Lai, and Chris Bates for helpful discussion. This research was supported by the Office of Naval Research (N00014-17-1-2984), the Center for Brains, Minds and Machines (funded by NSF STC award CCF-1231216), and a research fellowship from the Alfred P. Sloan Foundation.

## Footnotes

1 Selecting values larger than 0.1 resulted in some points lying above the reward-complexity curve, which is theoretically impossible and therefore indicates bias in the estimator. Different values of *a* do not significantly alter the shape of the empirical reward-complexity curve; the main effect is to shift the entire curve along the complexity axis.

2 Technically, compression will only be necessary if capacity constraint is lower than the number of bits needed to encode the optimal unconstrained policy. Here we are dealing with the case where the unconstrained policy is unachievable under the capacity constraint.

## Notes

### Competing Interest Statement

The authors have declared no competing interest.

